# A Hidden Proportional Feedback Mechanism Underlies Enhanced Dynamic Performance and Noise Rejection in Sensor-Based Antithetic Integral Control

**DOI:** 10.1101/2023.04.16.537062

**Authors:** Maurice Filo, Mucun Hou, M. Khammash

## Abstract

Efficient regulation of cellular processes is essential for both endogenous and synthetic biological processes. The design of biomolecular feedback controllers that achieve robust and timely regulation is the subject of considerable research at the interface between synthetic biology and control theory. Integral feedback controllers, known for their ability to confer the property of Robust Perfect Adaptation (RPA), are increasingly becoming common features in biological control design. Antithetic integral feedback (AIF) controllers, in particular, have enabled effective chemical reaction realizations of integral controllers that deliver RPA in both deterministic and stochastic settings. This paved the way to experimental implementations of integral controllers in bacterial and mammalian cells. While AIF controllers deliver favorable adaptation properties, they do not necessarily lead to good transient performance or noise reduction properties and may in some cases lead to increased overshoot or cell-to-cell variability. These limitations are commonly circumvented by augmenting new circuitry that realize proportional or derivative feedback mechanisms to enhance dynamic and noise rejection features without affecting the AIF controller’s adaptation properties. In this paper, we report that a sensor-based variant of the basic AIF motif exhibits favorable transient dynamic properties and (as reported elsewhere) reduced noise variance. We show that these features are attributed to a “hidden” proportional feedback component that is inherent in the controller structure and that such mechanism is solely responsible for the controller’s underlying enhanced dynamic performance and noise rejection properties. This sensor-based AIF controller hence offers a minimal biomolecular realization of Proportional-Integral (PI) control, whereby both integral and proportional feedback mechanisms are achieved through a single actuation reaction.

## 1 Introduction

Living cells are complex and dynamic systems that constantly interact with their environment. As a result, they are subject to various sources of disturbances, which can affect the accuracy and robustness of their biomolecular processes. To function properly, cells need to regulate these processes in a precise and timely manner, often relying on sophisticated feedback control mechanisms that ensure homeostasis. In recent years, the field of synthetic biology [1] has aimed to develop biomolecular circuits and systems that can be embedded inside the cells to mimic and even extend their natural control capabilities [2, 3]. These circuits and systems are built using genetic and biochemical components that can sense, process, and actuate signals in a programmable way. One of the key challenges in this field is to design and implement biomolecular feedback controllers that can handle noise and uncertainty, while achieving high performance and precision. The design of such synthetic controllers has been greatly influenced by new control-theoretic and engineering tools [4–8]. This led to the emergence of a new field of research at the interface of synthetic biology and control theory, referred to as Cybergenetics [9]. This intersection of disciplines has provided a fertile ground for the development of novel control strategies for biomolecular systems.

One of the fundamental tasks of synthetic biomolecular feedback controllers is to endow the overall control system with homeostasis. Controllers with such capabilities have direct implications and an unprecedented promising potential in various fields such as bioproduction, metabolic engineering, and especially in cell-based therapies due to the fact that many diseases are associated with loss of homeostasis [10]. A remarkable property known as Robust Perfect Adaptation (RPA) [11–13] is a special type of homeostasis that is more stringent than other types. It is a property which ensures the exact steady-state tracking of a variable of interest to a predetermined set-point despite varying initial conditions, uncertainties, and/or constant disturbances infiltrating the process to be regulated. According to the internal model principle [14], achieving RPA requires the controller to incorporate an integral feedback component. To this end, the antithetic integral feedback (AIF) controller [15] was introduced as a (bio)chemical reaction network that realizes integral feedback control and can be interconnected with an arbitrary network to achieve RPA in both the deterministic and stochastic settings. Later on, the AIF motif was shown to be not only sufficient to ensure RPA, but also necessary and minimal in the stochastic setting [16,17]. Although conceived from control-theoretic concepts, RPA-achieving controllers that leverage the AIF motif or its variants have rapidly found their way to experimental implementations in *Escherichia coli* [16, 18] and mammalian cells [19, 20].

Since the introduction of the AIF controller, efforts have been directed towards enhancing its performance either by tuning and exploring the dynamic trade-offs [21–23], or by adding extra circuitry [24–30] including molecular buffering, which was inspired by [31], and realizations of Proportional-Integral (PI) and Proportional-Integral-Derivative (PID) controllers. The need for such advanced biomolecular controllers arises from the limitations of standalone integral controllers. For example, previous works have highlighted that the standalone AIF controller can only shape the dynamic response up to a certain extent [24, 25]. Moreover, it was found that the standalone AIF controller achieves RPA at the population level, but it comes at the expense of increased stochastic noise in the form of elevated cell-to-cell variability [26]. However, these limitations can be mitigated by appending proportional and derivative components to the controller. In fact, studies have shown that adding a proportional feedback component to the AIF controller can improve dynamic performance and reduce the noise introduced by the integrator [24–26].

In this paper, we closely examine a simple variant of the AIF feedback motif which was first introduced in [15, Figure S1]. This variant is sensor-based and preserves the basic reaction network topology of the AIF network, but replaces a single actuation reaction with another one (see Fig. 2(c)). We reveal that this slight alteration actually gives rise to a PI controller that is realized with a single actuation reaction and without any additional circuitry. Despite being initially regarded as a standalone integral controller, we show that this variant in fact includes a “hidden” proportional component, which brings with it all the benefits of a proportional controller, such as enhancing the dynamic response and attenuating noise.

The paper is organized as follows. We begin by introducing the notation that we adopt throughout the paper. We then describe a general framework for biomolecular feedback controllers in Section 3. Afterwards, we describe three basic controller motifs in Section 4 that serve as the basis of the intuition behind the biomolecular filtered PI controllers designed and analyzed in Sections 5 and 6. We assess the deterministic dynamic performance of the AIF variant in Section 7. To do so, we employ standard analytical tools such as root locus and pole placements and back up the theoretical analysis with simulations. We then transition to the stochastic setting in Section 8 to explore the noise attenuation capabilities of the AIF variant. Finally we conclude with a discussion in Section 9.

## 2 Notation

Uppercase bold letters, e.g. **X**_**1**_, are reserved for species names. Their corresponding lowercase letters, e.g. *x*_1_(*t*), represent their deterministic concentrations as time-varying signals, whereas their corresponding uppercase letters, e.g. *X*_1_(*t*), represent their stochastic copy numbers, where *t* is time. Over-bars, e.g. 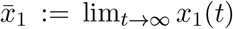, represent steady-state (or stationary) values when they exist. A tilde, e.g. 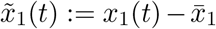, represents the deviation from the steady-state value, and a hat, e.g. 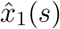, represents the Laplace transform of 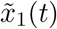, where *s* is the Laplace variable. The *s* and *t* variables are suppressed when they are clear from context. The Jacobian of a multi-variable function *f*, evaluated at 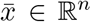 is denoted by 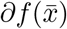. 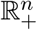 and 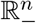 denote the set of *n*-sized vectors whose entries are non-negative and non-positive, respectively. *e*_*i*_ is a vector, of the appropriate size, whose entries are all zeros except the *i*^th^-entry being 1. Let 𝔼 [*X*_1_] and CV[*X*_1_] respectively denote the expectation and coefficient of variation of *X*_1_.

## 3 A Framework for Biomolecular Feedback Controllers

Consider the biomolecular closed-loop network depicted in Fig. 1(a) that describes a general framework for biomolecular feedback controllers adopted from [24]. It is comprised of an arbitrary network to be regulated, also referred to as the process or plant, interconnected in a feedback configuration with another network referred to as the controller network. The regulated network is comprised of *L* species: **X**_**1**_, **X**_**2**_, …, **X**_**L**_; whereas, the controller network is comprised of *M* species: **Z**_**1**_, **Z**_**2**_, …, **Z**_**M**_ reacting among each other. The two networks interact with one another through (1) a sensing reaction where the species **X**_**L**_, referred to as the regulated output, influences one (or more) controller species, and (2) an actuation reaction where one (or more) controller species influence species **X**_**1**_, referred to as the actuated input. Note that this framework describes a single-input/single-output (SISO) regulated network, but it can be straightforwardly generalized to the multiple-input/multiple-output (MIMO) case. The ultimate goal is to design a controller network that will ensure RPA such that the regulated output of interest **X**_**L**_ preserves a constant steady-state concentration despite the uncertainty and disturbances in the regulated network, and regardless of the initial conditions. In addition to RPA, the controller network needs to deliver a good dynamic performance and suppress undesirable noise.

**Figure 1:**
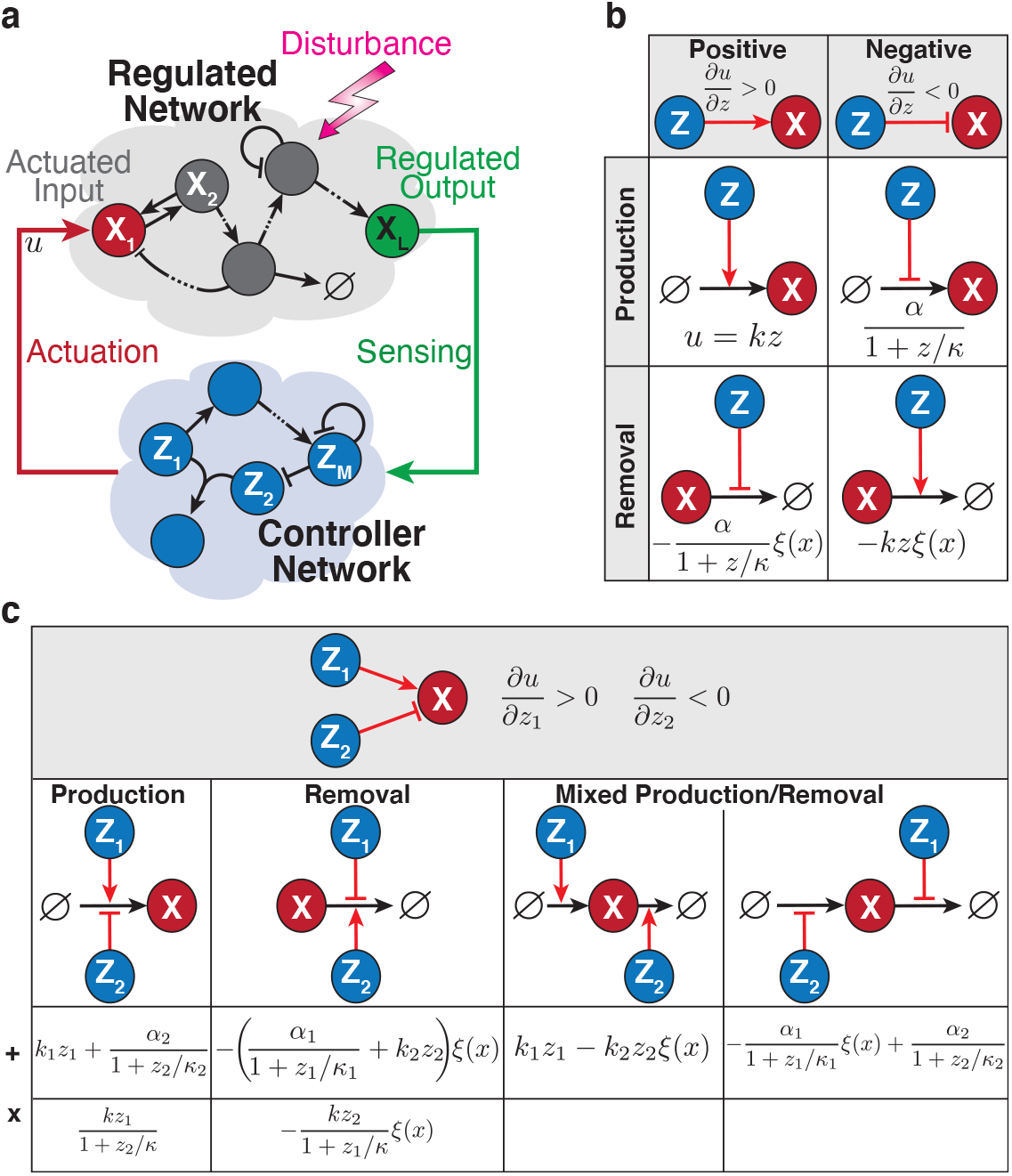
A framework for biomolecular feedback controllers. (a) Closed-loop biomolecular network. An arbitrary reaction network, referred to as the regulated network, is interconnected in a feedback configuration with the controller network whose objective is to endow the regulated output **X**_**L**_ with Robust Perfect Adaptation (RPA) – a property that provides robustness to disturbances and uncertainties – while ensuring high dynamic performance and reducing cell-to-cell variability. (b) Actuation mechanisms with a single controller species. Here, a single controller species **Z** actuates **X**. Positive actuation can be implemented as activating production or blocking removal; whereas negative actuation can be implemented as blocking production or activating removal. Mathematically, positive and negative actuations are captured by the sign of the derivative of the control action *u* with respect to *z*. Examples of the functional forms of *u* are provided. Note that *ξ*(*x*) denote the functional forms of degradation. (c) Actuation mechanisms with two controller species. This generalizes^**1**^ pannel (b) to the case where two controller species **Z**_**1**_ and **Z**_**2**_ actuate **X** positively and negatively, respectively. The implementations can once again be via production and/or removal reactions. Furthermore, two particular classes of functional forms are shown here, where the effects of **Z**_**1**_ and **Z**_**2**_ enter additively (such as separate promoters for the same gene) or multiplicatively (such as competition over the same promoter).

We consider the various actuation mechanisms tabulated in Fig. 1(b) which depicts actuations via one controller species, and Fig. 1(c), which depicts actuations via two controller species. In Fig. 1(b), the actuations are classified as either positive or negative and can be implemented via a production or removal reaction. More specifically, positive actuation can be accomplished by either increasing production or decreasing removal; conversely, negative actuation can be achieved by either reducing production or augmenting removal. As shown in Fig. 1(b), this can be represented by the sign of the derivatives of the control action *u* which can be computed as explained below. Consider the following actuation reactions with their propensities

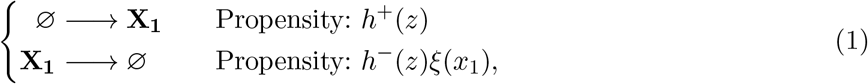

where the specific functional forms of *h*^*±*^ dictate the specific actuation mechanisms and *ξ*(*x*_1_) represents the functional form of degradation which can be linear or nonlinear to model degradation saturation, e.g. *ξ*(*x*_1_) = *x*_1_*/*(*x*_1_ + *κ*_*x*_). We define the total control action *u* as

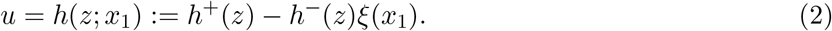

Various examples of the functional forms of *u* are provided in Fig. 1(b). In contrast, in Fig. 1(c), two controller species participate in the actuation such that **Z**_**1**_ actuates **X** positively, while **Z**_**2**_ actuates it negatively. Once again, these can be implemented via production and/or removal reactions. Finally, equipped with this framework, the dynamics of the closed-loop network in Fig. 1(a) can be generally written as

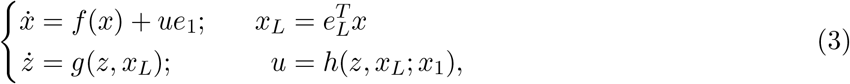

where (*f, g, h*) are some continuously differentiable functions, *x* := (*x*_1_ … *x*_*L*_]^*T*^ and *z* := (*z*_1_ *… z*]_*T*_. As a result, the feedback control problem boils down to the design of the functions *g* and *h* to ensure RPA while simultaneously delivering a high dynamic performance and possibly suppressing noise.

## 4 Biomolecular Proportional & Feedforward Controllers

Consider the three basic biomolecular controller topologies depicted in Fig. 2(a). The dynamics for all of the three basic controllers can be compactly written as

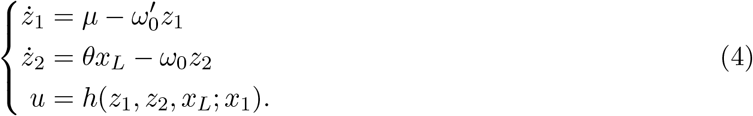

The dynamics of each of the three controllers can be obtained by suitably choosing the actuation function *h*. To unravel the underlying controller architectures, we carry out a standard linear perturbation analysis. The derivation details are straightforward and can be found in Appendix A.1. The analysis yields the controller transfer function of the linearized perturbation dynamics from *x*_*L*_ to *u* given as

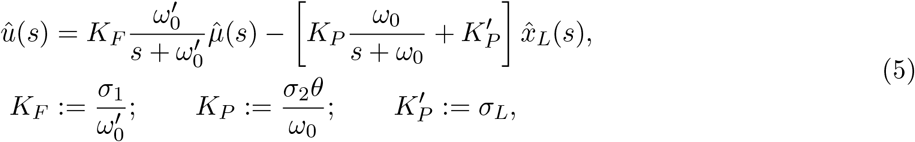

where the partial derivatives of the actuation function *h* are defined as

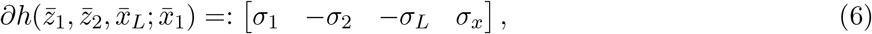

with *σ*_1_, *σ*_2_, *σ*_*L*_ ≥ 0. The controller transfer function in (5) allows us to draw the block diagram depicted in Fig. 2(b). This block diagram unravels the architectures of the three controllers. The proportional controller delivers an instantaneous feedback from the output **X**_**L**_ to the input **X**_**1**_. The filtered proportional controller, on the other hand, passes the proportional control action through a low-pass filter which is realized as a simple birth-death process via an intermediate species **Z**_**2**_ between the output **X**_**L**_ and input **X**_**1**_. Finally, the feedforward control action has no feedback from the output **X**_**L**_, but it is also passed through a low-pass filter.

**Figure 2:**
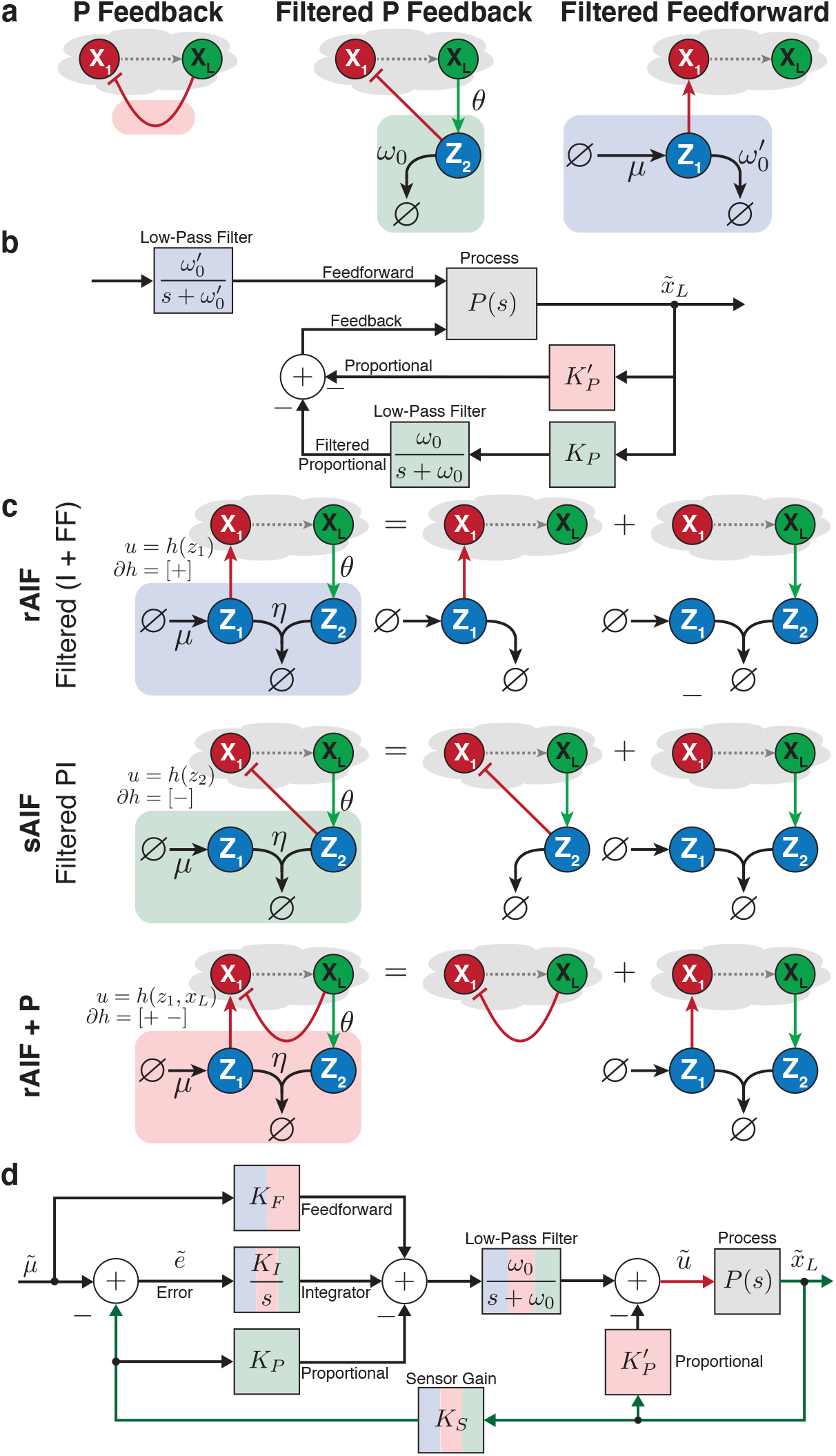
Topological Assembly of biomolecular PI controllers. (a) Basic biomolecular controller motifs. The topology to the left depicts a direct negative feed-back control action that doesn’t pass through any intermediate controller species. In contrast, the topology in the middle introduces an intermediate controller species **Z**_**2**_ whose production is driven by the regulated output **X**_**L**_, degrades at a rate *ω*_0_ and negatively actuates the input **X**_**1**_. The topology to the right involves an intermediate controller species **Z**_**1**_ which is constitutively produced at a rate *μ*, degrades at a rate 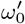 and positively actuates **X**_**1**_. (b) The underlying control architectures of the basic controller motifs. *P* (*s*) denotes the transfer function of the process, and the other blocks are color-coded to match the three basic controller motifs. Linear perturbation analysis reveals that instantaneous feedback realizes a “pure” proportional controller with gain 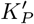; while the intermediate species appends to the proportional controller a low-pass filter whose cutoff frequency is *ω*_0_. In contrast, the motif to the right embeds a feedforward controller (excluding any feedback) with a low-pass filter whose cutoff frequency is 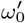. (c) and (d) Effects of combining a sequestration motif with the three basic controller motifs. The distinction between the topologies of the reference-based and sensor-based AIF controllers lies in the actuation reaction: for rAIF, the reference molecule **Z**_**1**_ positively actuates **X**_**1**_; whereas for sAIF, the sensor molecule **Z**_**2**_ negatively actuates **X**_**1**_. This subtle difference results in entirely distinct control architectures, where rAIF appends the integrator with a feedforward component with gain *KF* ; while sAIF appends it with a proportional component with gain *KP*. The resulting PI architecture is hence achieved through a single actuation reaction, as opposed to the separate Proportional (with gain 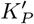) and Integral components realized in the last topology (rAIF + P).

## 5 Biomolecular Proportional-Integral Controllers

In this section, we “append” the basic controller motifs listed in Fig. 2(a) to the sequestration motif – which lies at the heart of the antithetic integral feedback (AIF) controller [15] – to obtain the three topologies depicted in Fig. 2(c). Afterwards, we examine the resulting underlying controller architectures.

To obtain the reference-based (rAIF) and sensor-based (sAIF) controllers, we append the sequestration motif, to the filtered feedforward component and the filtered proportional component, given in Fig. 2(a), respectively. Whereas to obtain the topology in the last row of Fig. 2(c), we append the rAIF controller with the pure (non-filtered) proportional component. Once again, using a standard linear perturbation analysis (see Appendix A.2 for the derivation details), we unravel the control architectures that are compactly summarized in a single block diagram depicted in Fig. 2(d). Note that the block diagram is color-coded consistently with the controller reaction networks to highlight the blocks corresponding to each of the three topologies. The analysis reveals that rAIF realizes integral and feedforward control that are both passed through a low-pass filter, whereas sAIF realizes a PI controller that is passed through a low-pass filter. Finally, the last topology in Fig. 2(c) has the same control architecture as the rAIF topology, but with an additional pure proportional component.

To conduct a simulation-free evaluation of the dynamic capabilities of these controller topologies, we examine the achievable ranges of the gains (*K*_*P*_, *K*_*I*_, …) and cutoff frequency *ω*_0_. Specifically, we ask: can we adjust these gains and cutoff frequency to any desired value? If not, what are the achievable values? Of course, a broader achievable range indicates greater flexibility in shaping the dynamic response. To address these questions in a simulation-free manner, we first establish a bi-directional mapping between the biomolecular parameters and the gain/cutoff-frequency parameters. This enables us to map the constraints on biological parameters (such as positivity) into the gain/cutoff-frequency parameter space, thereby revealing the attainable range. In the following section, we conduct this analysis for the sAIF topology. The analysis for the rAIF topology can be found in Appendix A.2, while that for the rAIF + P topology is available in [24, 25].

## 6 Mappings between the Filtered PI and Biomolecular Parameter Spaces

Consider the sAIF controller depicted in Fig. 2(c), where **Z**_**2**_ negatively actuates the input species **X**_**1**_, that is, the actuation function *h*(.) is monotonically decreasing in *z*_2_. Recall that we established in the previous section that this topology realizes a filtered PI controller. Our goal is to derive the mappings between the Filtered PI parameters (*K*_*P*_, *K*_*I*_, *ω*_0_) and the various biomolecular parameters (*η*, …), noting that for sAIF, 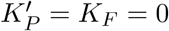. We first start with the analysis problem: given the biomolecular parameters, what are the PI gains and cutoff frequency? Then we move to the design problem: what are the biomolecular parameters that achieve some desired PI gains and cutoff frequency?

Throughout the subsequent analysis, we will make an assumption about the process. Let *F*_*i*_ (*i* = 1, 2, …, *L*) denote the steady-state maps of the process, that is, if *u* is a constant then with reference to the first equation in (3), we write

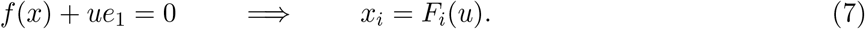

### Assumption 1

*Assume that for the desired steady-state output* 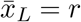, *there exists a feasible supporting input ū and steady-state concentrations of the process species* 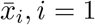, *i* = 1, …, *L* − 1, *that achieve the desired output. More precisely, r >* 0, ∃*ū* ∈ U, 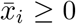 *such that F*_*L*_(*ū*) = *r and* 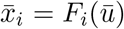, *where 𝕌 is the set of feasible inputs*.

### Remark 1

*The set of feasible inputs depends on the type of actuation. For instance if the actuation is carried out via non-saturating production only, then 𝕌* = ℝ_+_; *whereas if it is carried out via non-saturating degradation only, then 𝕌* = ℝ_−_. *If both non-saturating production and degradation actuations are allowed then, 𝕌* = ℝ.

### Remark 2

*We emphasize that this assumption does not depend on the type of controller used. Instead, it only depends on the process and the particular choice of actuated input species and actuation mechanism. This assumption has to be satisfied, otherwise, the actuation is simply inadequate and there is no controller that can achieve the desired output without changing the choice of the actuated input species and/or actuation mechanism*.

Next, we treat the analysis and design problems for two biologically-relevant functional forms of *h* implementing the two negative actuation mechanisms (production and removal) shown in Fig. 1(b).

### 6.1 Repression

The actuation function *h* is given here as a Hill-type function with cooperativity, that is

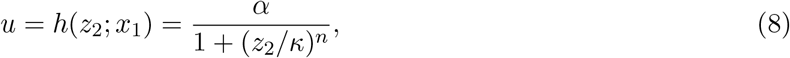

where *κ* is the dissociation constant, *α* is the maximal production rate and *n* is the Hill coefficient. The set-point is given by 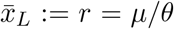. For a given process, satisfying Assumption 1, and set-point *r*, the supporting input *ū* satisfies *F*_*L*_(*ū*) = *r* and is fixed. We first treat the analysis problem, then move on to the design problem.

#### Analysis

The controller coordinates 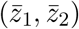 of the fixed point are given by

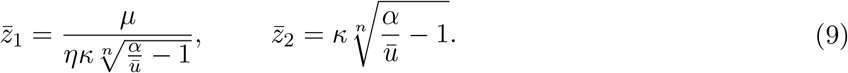

Clearly, the following condition on the biomolecular parameters has to be satisified to guarantee that 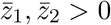,

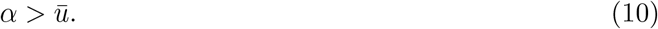

Violating this condition causes both coordinates of the fixed point to become either negative or complex and thus causing instability. By substituting the partial derivatives of the actuation function *σ*_1_ = *σ*_*L*_ = *σ*_*x*_ = 0 in (30), one can write the PI gains (*K*_*P*_, *K*_*I*_) and cutoff frequency *ω*_0_ in terms of the various biochemical parameters as

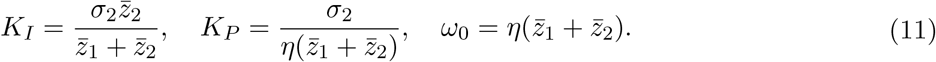

where 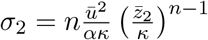.

#### Design

By fixing *μ* and *r* (and thus *ū*), one can easily solve the equations given in (9) and (11) for the biomolecular parameters *α, κ*, and *η* in terms of the PI gains and cutoff frequency to obtain

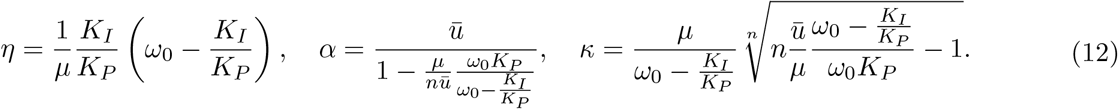

Since the biomolecular parameters *α, κ* and *η* cannot be negative and have to satisfy the condition in (10), the achievable PI gains and cutoff frequency are constrained as described next.

#### Filtered-PI Coverage

Constraining *α, κ* and *η* to be non-negative and to satisfy condition (10) yields the following achievable PI gains and cutoff frequency.

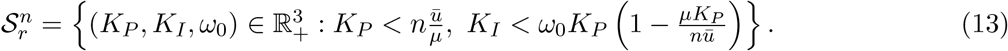

This indicates that employing repression for negative actuation imposes an upper bound on both the proportional gain *K*_*P*_ and integral gain *K*_*I*_. It is worth noting that these upper bounds can be relaxed by increasing *n* since 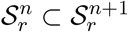, suggesting that cooperativity enhances the coverage, thereby enabling more flexible tuning of the filtered PI parameters. Lastly, it is important to highlight that the upper bound of *K*_*I*_ depends not only on the process and the set-point via the supporting input *ū*, but also on the proportional gain *K*_*P*_ and cutoff frequency *ω*_0_.

### 6.2 Degradation

Next, consider the case where **Z**_**2**_ degrades the input species **X**_**1**_. The actuation function *h* is thus given by

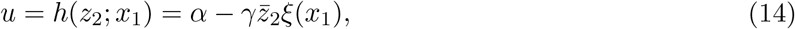

where 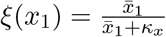. The controller coordinates 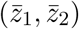 of the fixed point are

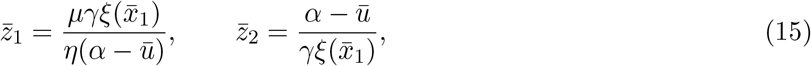

with 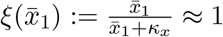, by choosing *κ*_*x*_ to be small for simplicity. Note that this assumption can be easily relaxed. Calculations of the analysis problem are similar to the repression case but with 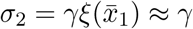. The mapping from the PI gains (*K*_*P*_, *K*_*I*_) and the cutoff frequency *ω*_0_ to the biomolecular parameters is

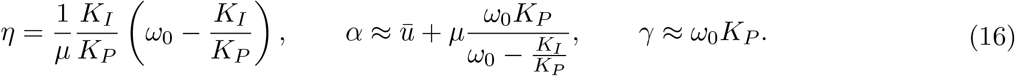

Constraining the biomolecular parameters to be non-negative yields the following achievable PI parameters,

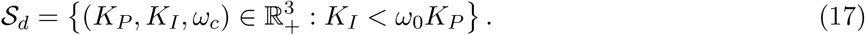

This indicates that employing degradation for negative actuation, imposes an upper bound on the integral gain *K*_*I*_ only. Furthermore, this bound is less restrictive than that corresponding to the actuation via repression. In fact, observe that for all *n* = 1, 2, …, we have 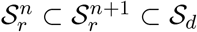 as visually demonstrated in Fig. 3. It is worth mentioning that this filtered PI controller has more constrained coverage compared to the filtered and non-filtered PI controller in two actuation reactions [25,32]. This is the price of having a minimal design where both the proportional and integral components are realized in a single actuation reaction.

**Figure 3:**
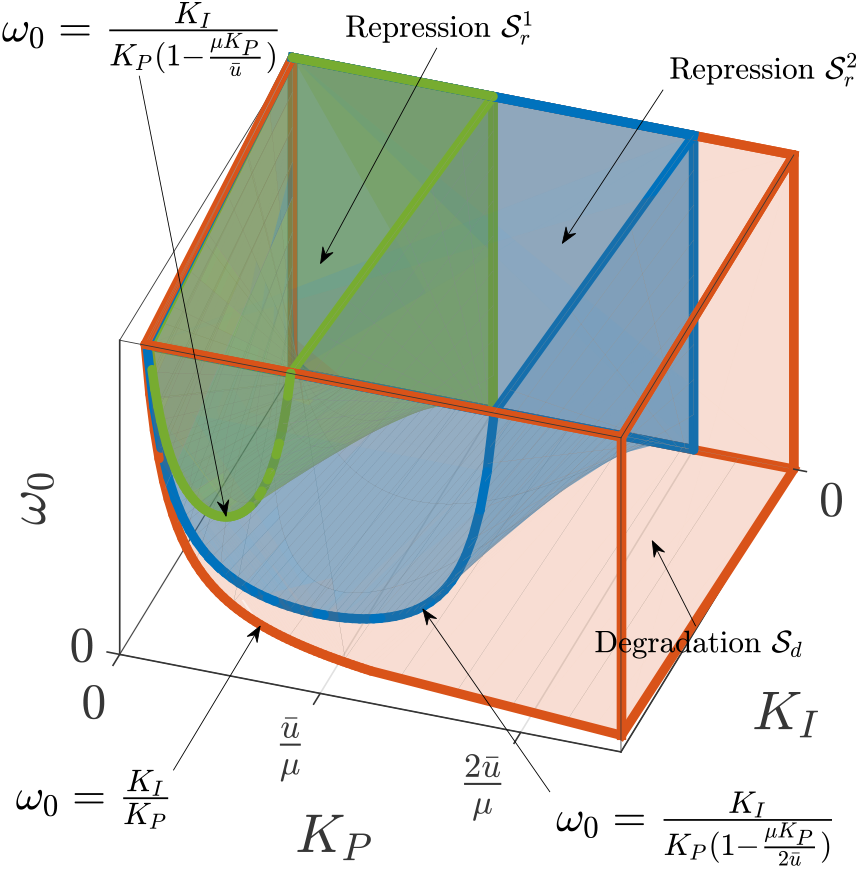
Filtered-PI Coverage. The colored regions depict the achievable PI gains (*K*_*P*_, *K*_*I*_) and cutoff frequency *ω*_0_ by adjusting the corresponding biomolecular parameters. These regions are color-coded to represent different actuation functions *h*, modeling three distinct negative actuation mechanisms: repression (8) with and without cooperativity in green (*n* = 1) and blue (*n* = 2), respectively, and degradation (14) in red. Note that *ū* represents the steady-state supporting input necessary to achieve the desired set-point, and its value depends solely on the plant and the desired set-point. The span of achievable filtered-PI parameters for repression and degradation actuations are respectively calculated as 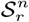 in (13) and **𝒮**_*d*_ in (17), and they are shown to satisfy 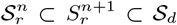. This demonstrates that degradation provides greater tuning flexibility than repression inhibition. It also demonstrates that cooperativity helps in expanding the achievable gains and cutoff frequency.

## 7 Dynamic Performance Assessment

The objective of this section is to demonstrate, analytically and through simulations, that negative actuation by **Z**_**2**_ (sAIF controller in Fig. 2(c)), enables more flexibility in enhancing the dynamic performance when compared to positive actuation by **Z**_**1**_ (rAIF controller in Fig. 2(c)). Furthermore, we also explore the performance-enhancement capabilities of the two negative actuation mechanisms presented in Section 6. This superior performance is a direct consequence of the additional filtered proportional component that is not present when actuating with **Z**_**1**_.

Consider the closed-loop dynamics with a one-species (*L* = 1) birth-death process, for simplicity. That is *f* (*x*) = −*γ*_1_*x* + *u*. This simple plant is enough to demonstrate the added flexibility of the proportional component. The transfer function of this plant is given by 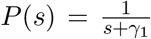. Using the block diagram in Fig. 2(d), it is straightforward to write down the closed-loop transfer functions of the linearized dynamics for rAIF (*K*_*P*_ = 0, *K*_*F*_ *>* 0) and sAIF 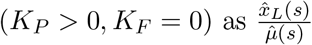:

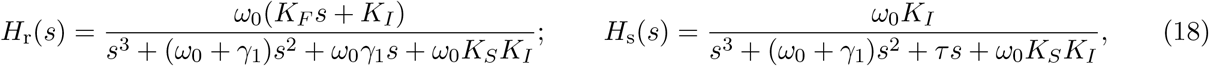

where *τ* := *ω*_0_(*γ*_1_ + *K*_*P*_ *K*_*S*_). Using simple root-locus arguments (see Appendix B), it can be shown that for *H*_*r*_(*s*), at least one pole cannot be placed to the left of 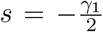 no matter how we tune *K*_*I*_ and even if we resort to a fast cutoff frequency *ω*_0_ (that is fast sequestration rate *η*). This means that the performance of the rAIF topology realizing a filtered (I+FF) controller is limited in the sense that the speed of the response cannot exceed a threshold dictated by 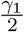. This is analytically established in Appendix B by deriving an analytical formula for the breaking point *s*_*b*_ of the root locus given in (36) and visually demonstrated in Fig. 4(a).

**Figure 4:**
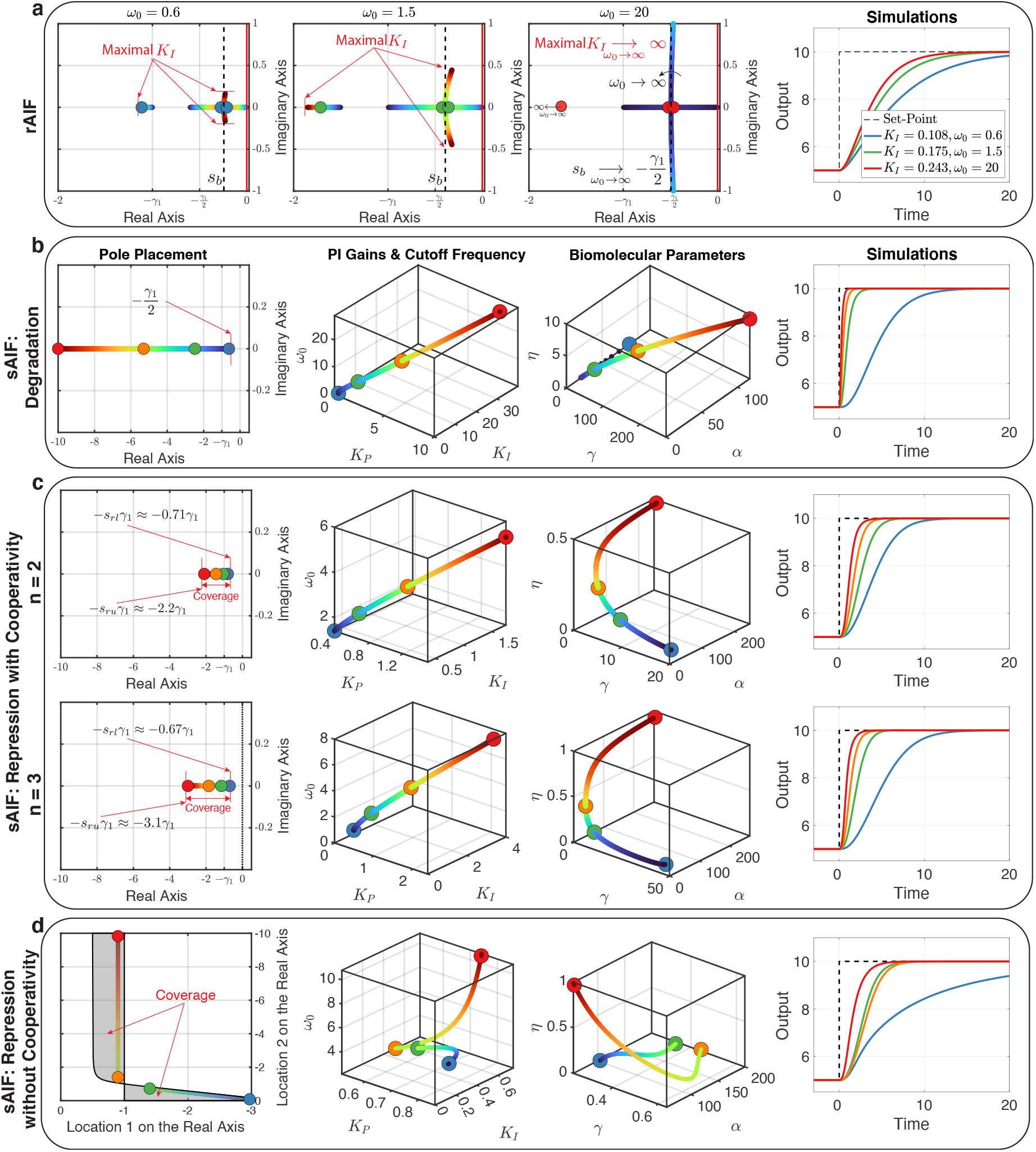
Dynamic Performance Assessment. A birth-death process (see Fig. 5(a), left) is controlled, as a case study, by rAIF and sAIF. The control action is denoted by *u* and the degradation rate of the process is denoted by *γ*_1_. (a) Performance limitation of rAIF. Positive actuation by **Z**_**1**_ (i.e. *u* = *kz*_1_) yields a response that cannot be sped up beyond a certain threshold without inflicting oscillations. The three plots to the left depict the root locus of the linearized closed-loop dynamics in the complex plane for three values of the cutoff frequency *ω*_0_ as the integral gain *K*_*I*_ is increased from zero up to its upper bound given in (33). Note that *s*_*b*_, calculated analytically in (36), denotes the breaking point where two eigenvalues meet on the real axis and break away to become complex conjugates. As *ω*_0_ is increased, one real eigenvalue moves more to the left and the breaking point *s*_*b*_ tends to −*γ*_1_*/*2. This indicates that the dominant eigenvalue is confined (by the breaking point *s*_*b*_) within a small region close to the imaginary axis when *γ*_1_ is small, and thus imposing a limitation on the achievable performance as demonstrated in the simulations shown in the right plot. (b) and (c) Design flexibility offered by sAIF. Giving rise to a filtered-PI controller, sAIF offers more flexibility in achieving superior performance compared to rAIF. These two panels show the steps of a pole-placement control design problem where the three dominant poles are placed on the real axis of the left-half plane to ensure a stable and non-oscillating response. The design problems start by picking the poles, then computing the PI gains and cutoff frequency, and finally computing the actual biomolecular parameters that allow us to obtain the nonlinear simulations to the right. With degradation actuation in Panel (b), one can place the eigenvalues arbitrarily as far to the left as desired and thus achieving a response that is as fast as desired without overshoots or oscillations. In contrast, with repression in Panel(c), there is a restriction on how far to the left the poles can be placed. However, this restriction can be mitigated by introducing higher cooperativity. (d) Repression without cooperativity. Without cooperativity, the three poles cannot be placed in the same location. To this end we choose to place them at two locations on the real axis. The shaded regions in the plot to the left depicts the feasible locations that are constrained by the PI coverages (see Appendix C). These regions indicate that one cannot place all the poles to the left of −*γ*_1_ which still yields a better performance than rAIF, but cannot outperform those presented in Panels (b) and (c). The numerical values of the fixed parameters are *γ*_1_ = 1, *μ* = 5, *θ* = 1, *κ*_1_ = 10^−5^. To change the set-point at *t* = 0, *μ* is doubled.

Here is exactly where the filtered-proportional component, arising from the actuation via **Z**_**2**_ instead of **Z**_**1**_, comes into play to add more flexibility. To showcase this flexibility, let’s consider the pole placement design problem. The objective is to design the PI gains (*K*_*P*_, *K*_*I*_) and cutoff frequency *ω*_0_ in a way that places the three closed-loop poles at a specific location *s* = −*a*. Ideally, this location should be in the left-half complex plane to ensure stability and on the real axis to prevent oscillations. Moreover, placing it far to the left will result in a fast transient response, which is desirable. These conditions can be met by selecting a sufficiently large value of *a >* 0. We will now investigate whether this can be achieved with sAIF using the two negative actuation mechanisms: repression and degradation. If this is achievable, we examine the pole placement range; otherwise, we aim at placing the poles at two distinct locations and analyze their effect on the dynamics.

First, we aim at placing the three closed-loop poles at the same location *s* = −*a*. As a result, the characteristic polynomial is given by

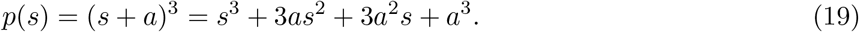

Equating *p*(*s*) to the denominator of *H*_s_(*s*) allows us to express the designed PI gains (*K*_*P*_, *K*_*I*_) and cutoff frequency *ω*_0_ in terms of the birth-death parameter *γ*_1_, the sensing gain *K*_*S*_ and the placed pole −*a* as

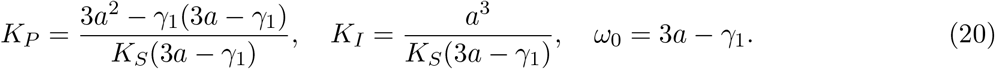

The sets of achievable PI gains and cutoff frequencies (Fig. 3) constrain the achievable poles *s* = −*a* to the following regions on the real axis.

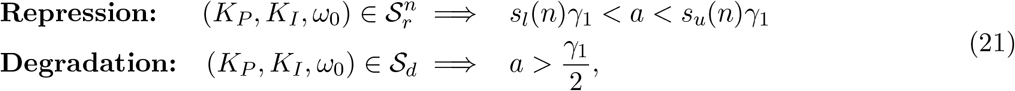

where *s*_*l*_(*n*) and *s*_*u*_(*n*) are calculated analytically in Appendix C. Observe that with degradation actuation, there is no theoretical upper limit. This is numerically demonstrated in Fig. 4(b) where the poles are placed as far to the left as desired to yield an ideal step-like response exhibiting no overshoots, no oscillations and fast transient. This demonstrates that sAIF is capable of freely shaping the dynamic response of a birth-death process while rAIF fails to do so. This is due to the additional embedded proportional component that is achieved by the negative actuation with **Z**_**2**_ instead of the positive actuation with **Z**_**1**_. In contrast, when actuating via repression, the pole locations are constrained to be in the open set **ℛ** (*n*) = (−*s*_*u*_(*n*)*γ*_1_, −*s*_*l*_(*n*)*γ*_1_). In fact, without cooperativity (*n* = 1), we have *s*_*l*_(1) = *s*_*u*_(1) = 1 which means that **ℛ** (1) is empty. To this end, the poles cannot be placed at the same location on the real axis. However, for *n* = 2, *s*_*l*_(2) ≈ 0.7082 and *s*_*u*_(2) ≈ 2.1769. This indicates that cooperativity is necessary to place the poles at the same location on the real axis. Additionally, the range **ℛ** (*n*) of achievable poles is broadened in the presence of a higher degree of cooperativity (larger *n*) as demonstrated in Fig. 4(c).

To understand the dynamical capability of sAIF when the actuation is done via a repression reaction without cooperativity (i.e. *n* = 1), we aim at placing the three poles at two different locations (two poles at *s* = −*a*_1_ and one pole at *s* = −*a*_2_) since placing them at a single location is not possible.

The coverage of the gains and cutoff frequency (*K*_*P*_, *K*_*I*_, *ω*_0_) constrains the poles in a bounded region which is calculated in Appendix C. This region indicates that the two locations cannot be chosen to the left of *s* = −*γ*_1_ simultaneously, as numerically demonstrated in Fig. 4(d). Consequently, the speed of the transient response is limited by a threshold dictated by *γ*_1_. Note that this limitation is still less constrained compared to rAIF where the threshold is dictated by 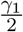.

In conclusion, this case study (controlling a birth-death process) shows that sAIF delivers superior performance compared to rAIF. In particular, implementing the negative actuation of sAIF via a degradation reaction provides enough degrees of freedom to arbitrarily speed up the transient response without giving rise to any overshoots or oscillations. In contrast, although implementing the negative actuation of sAIF via repression also improves the performance compared to that of rAIF, the responses cannot be made arbitrarily fast without incurring oscillations and/or overshoots. This performance limitation, however, can be mitigated by introducing cooperativity in the repression.

## 8 Limits of Stochastic Noise Attenuation

In this section, we transition to the stochastic setting and conduct a simulation study to explore the noise attenuation capabilities of three controllers: rAIF, sAIF and filtered proportional controllers (refer to Fig.2). We begin by defining stochastic noise as the relationship between the coefficient of variation and the expectation at stationarity. We consider two processes illustrated in Fig.5(a), representing birth-death and gene expression models. For the birth-death process, the output of interest is **X**_**1**_, while for the gene expression process, the output of interest is **X**_**2**_. Throughout the analysis, the parameters of the processes (i.e. *γ*_1_, *γ*_2_, and *k*_1_) are fixed. The two processes are controlled by one of the three controllers depicted in Fig. 5(a). Note that negative actuations are implemented via a repression reaction. In the open-loop scenarios, the control action *u* = *α* is a constant rate. This gives rise to unimolecular networks with closed moment equations. In fact, the coefficients of variation for the outputs at stationarity can be explicitly expressed as functions of their expectations:

**Figure 5:**
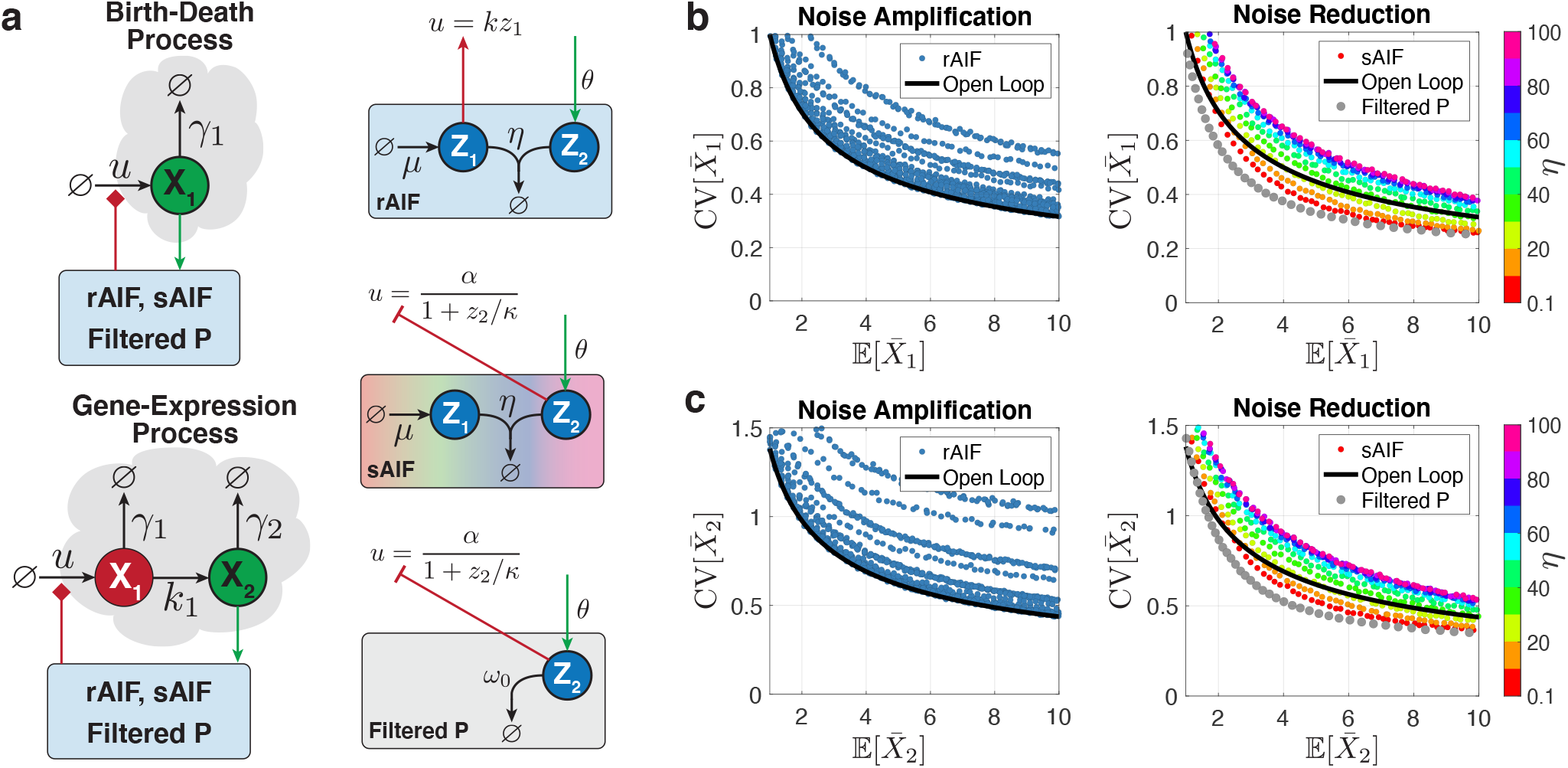
Stochastic noise attenuation capabilities and limitations. (a) We examine two case studies for the processes to be regulated: a birth-death process and a gene expression process. These processes are controlled by three different controllers: rAIF and sAIF, which supplement integral controllers with filtered feedforward and proportional components, respectively, and a filtered proportional controller without an integrator. (b) and (c) display the relationship between the coefficients of variation and expectations at stationarity for the birth-death and gene expression processes, respectively. The left plots correspond to rAIF, while the right plots correspond to sAIF and filtered proportional feedback. The key takeaway from these plots is that rAIF can only increase noise compared to the open-loop scenario, while sAIF can attenuate noise to a certain extent, limited by its “hidden” proportional component. The simulations support the notion that integral controllers amplify noise, whereas proportional controllers attenuate it. The solid black lines are calculated analytically using (22), while the various circles are computed empirically through the stochastic simulation algorithm [33], generating 10^4^ − 10^5^ trajectories on the Euler cluster (https://scicomp.ethz.ch/wiki/Euler). Numerical values for the birth-death process are: *γ*_1_ = 0.1. For the gene expression model, parameter values are: *γ*_1_ = *k*_1_ = 1, *γ*_2_ = 0.1. The controller parameter values are as follows: *α* = 2, *θ* = 1, *κ* = 0.05, *η* ∈ [0.1, 100], *k* ∈ [10^−3^, 1], *ω*_0_ ∈ [10^−2^, 10^3^], *μ* ∈ [1, 10].

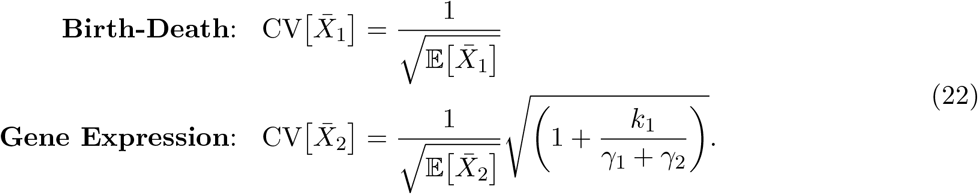

These analytical expressions are represented as solid black curves in Fig.5(b) and (c).

Next, we carry out stochastic simulations for the closed-loop scenarios using each of the three different controllers. These simulations compute the stationary expectations and coefficients of variation for the outputs across a range of specific controller parameters. In particular, for rAIF, we vary *k, η*, and *μ*, while keeping *θ* constant. The results for each process are shown as data points in Fig.5(b) and (c) on the left. The simulation findings indicate that rAIF can only increase noise compared to the uncontrolled open-loop scenario. For sAIF, we vary *η* and *μ* while keeping *α, κ*, and *θ* constant. The results for each process are displayed as color-coded data points, for different *η* values, in Fig.5(b) and (c) on the right. In contrast to rAIF, the simulations reveal that sAIF can reduce noise to levels below those observed in the uncontrolled open-loop setting. For the filtered proportional controller, comparable stochastic simulations are conducted for different values of *ω*_0_. Note that, in order to perform a fair comparison, we fix the same parameters as those of the sAIF controller. The results are displayed as gray data points in Fig.5(b) and (c) on the right, indicating that the limit of noise reduction delivered by sAIF is dictated by the filtered proportional controller. This suggests that the noise attenuation in sAIF is attributable to the “hidden” proportional component, rather than the integrator. In fact, as *η* increases in the sAIF controller, noise also increases, which is consistent with the proportional gain *K*_*P*_ in (11) approaching zero as *η* approaches infinity.

## 9 Discussion

Achieving Robust Perfect Adaptation (RPA) through biomolecular controllers is crucial in regulating cellular processes in living cells, which are inherently noisy and uncertain. While RPA is a necessary property to maintain cellular homeostasis, it is often not sufficient for achieving high performance. Furthermore, RPA may be achieved at the population level but at the expense of high cell-to-cell variability. Therefore, there is a need to develop biomolecular controllers that can deliver both RPA and high performance, taking into account the inherent variability of living cells.

While integral controllers are usually the suitable choice to achieve RPA at the population level [15], proportional controllers are often added on top of the integrators to enhance the dynamic performance and reduce noise or cell-to-cell variability [24,26]. In previous works, such addition was usually realized by adding extra circuitry which could be biologically demanding, although unavoidable in certain scenarios. In this paper, we have shown that a slight variant of the standard rAIF controller (see the sAIF topology in Fig. 2(c)) gives rise to a (filtered) PI controller without adding the extra circuitry. We also demonstrated analytically and through simulations that this variant indeed brings in the benefits of the proportional controller while maintaining the RPA property offered by the integrator.

The sAIF controller was first introduced in [15, Fig. S1] as one of several realizations of AIF control. More recently, a stochastic analysis employing linear noise approximation was conducted in [34] to show that this variant is capable of reducing noise when controlling a birth-death process. Our study reveals that it is precisely the “hidden” proportional component which is responsible for this noise reduction, and not the integrator. This is demonstrated in Fig. 5 when regulating not only a birth-death process but also a gene expression process. We also demonstrate analytically and through simulations that the “hidden” proportional component not only reduce noise, but also enhances the dynamic performance. Interestingly, this seemingly minor, but subtle, alteration in the choice of the actuating species yields a different controller architecture which tangibly offers better responses. The intuition behind this improvement lies in the fact that the altered choice of actuating species cascades both a filtered proportional controller and an integral controller, resulting in the best of both worlds. This finding has practical implications for biologists as it offers a minimal design for biomolecular PI controllers which is easier to build.

## A Transfer Functions

Let 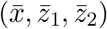 denote the closed-loop fixed point when operating at a nominal exogenous input 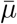. Furthermore, let 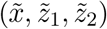 denote the perturbation from the closed-loop fixed point due to a disturbance or a perturbation 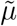 of the exogenous input from its nominal value. That is, we have

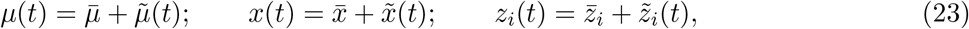

for *i* = 1, 2. Let 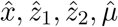 and *û* respectively denote the Laplace transforms of 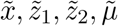 and *ũ*. For the actuation function *h*, define the partial derivatives as 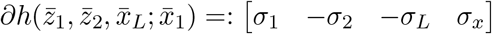 with *σ*_1_, *σ*_2_, *σ*_*L*_ ≥ 0, and let *e*_*i*_ be a vector of an appropriate size whose entries are all zeros except the *i*^th^-entry being 1.

### A.1 Proportional & Feedforward Controllers

Consider the following closed-loop dynamics

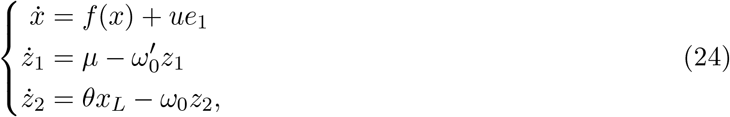

where the control action is given as *u* = *h*(*z*_1_, *z*_2_, *x*_*L*_; *x*_1_) to encompass all three basic controller motifs listed in Fig. 2(a) and discussed in Section 4. The approximated perturbation dynamics are thus given by the linearization that can be written separately for the process **𝒫** and the controller **𝒞** as

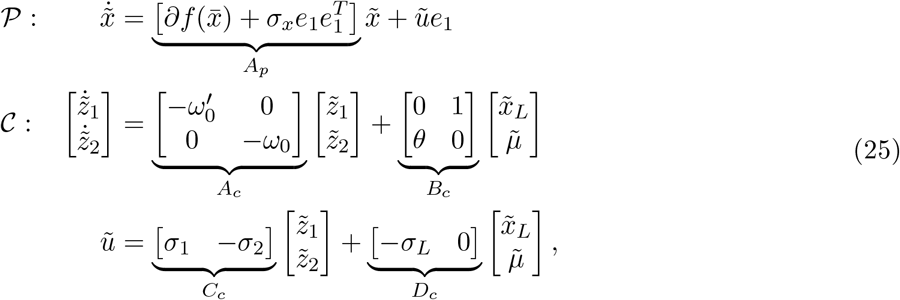

where, for convenience and with a slight abuse of notation, *σ*_*x*_ is absorbed in the dynamics of the process and so 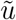 does not involve 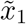. Taking the Laplace transforms on both sides of the equalities in (25) and recalling that the transfer matrix of the controller is *C*_*c*_(*sI* − *A*_*c*_)^−1^*B*_*c*_ + *D*_*c*_ yields the following transfer functions

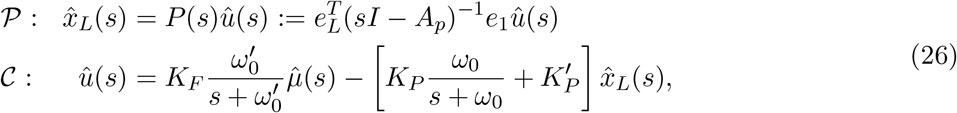

where

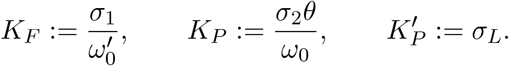

As a result, the three cases presented in Fig. 2(a) can be directly obtained from (26) by choosing the control action *u* = *h*(*z*_1_, *z*_2_, *x*_*L*_; *x*_1_) appropriately which leads to setting a subset of the partial derivatives *σ*_1_, *σ*_2_, *σ*_*L*_ and *σ*_*x*_ to zero.

### A.2 Proportional-Integral Controllers

Consider the following closed-loop dynamics

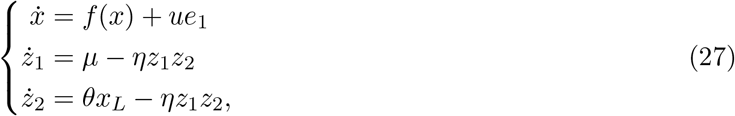

where the control action is given as *u* = *h*(*z*_1_, *z*_2_, *x*_*L*_; *x*_1_) to encompass all three cases presented in Fig. 2(c) and (d) which are discussed in Section 5. The approximated perturbation dynamics are thus given by the linearization that can be written separately for the process **𝒫** and the controller **𝒞** as

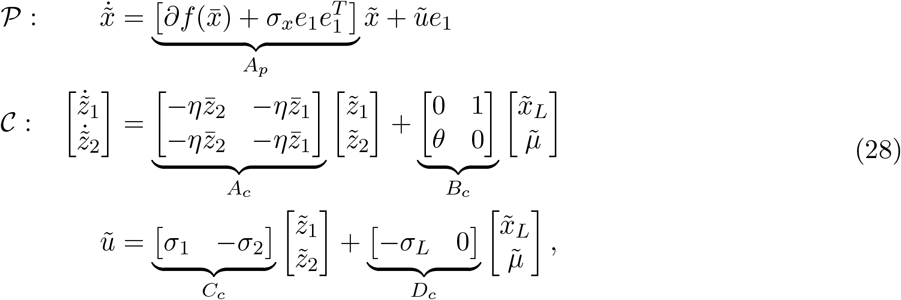

where, for convenience and with a slight abuse of notation, *σ*_*x*_ is absorbed in the dynamics of the process and so *ũ* does not involve 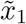. Taking the Laplace transforms on both sides of the equalities in (28) and recalling that the transfer matrix of the controller is *C*_*c*_(*sI* − *A*_*c*_)^−1^*B*_*c*_ + *D*_*c*_ yields the following transfer functions

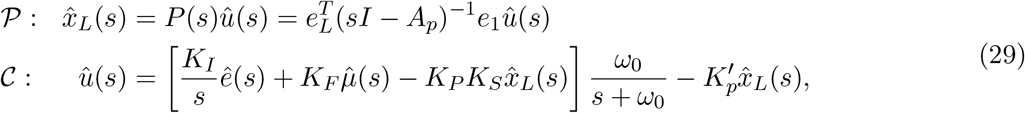

where

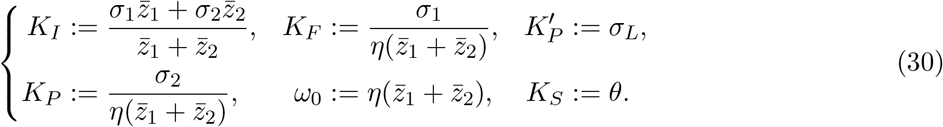

This set of formulas provides a way to calculate the block diagram parameters (see Fig. 2(d)) from the biomolecular parameters. To go in the opposite direction, one can solve (30) for the biomolecular parameters to obtain

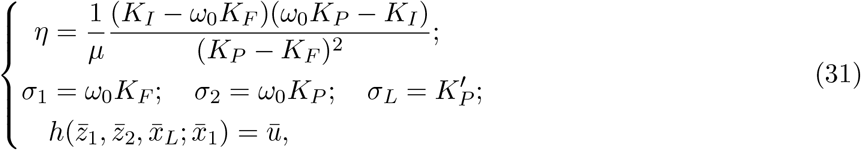

where 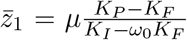 and 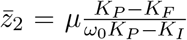. Of course whether this inversion is doable or not depends on the number of degrees of freedom that shape the actuation function *h*. In particular for rAIF with *h*(*z*_1_, *z*_2_, *x*_*L*_; *x*_1_) = *kz*_1_, we have 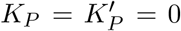 and for a fixed *μ* and *θ*, the mappings back and forth between the block diagram and biomolecular parameters are given by

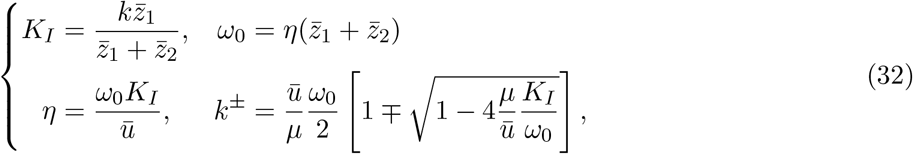

Observe that *K*_*F*_ is left out on purpose because with this actuation function *K*_*F*_ is not a degree of freedom (unless *μ* or *θ* are allowed to be tuned). Furthermore, since *k* has to be a nonnegative real number, then the following condition constrains the coverage of the integral gain and cutoff frequency:

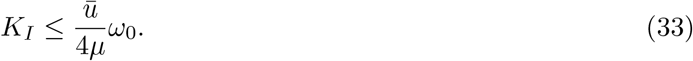

For sAIF, we have 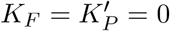 since *h* is not a function of *z*_1_. The mappings back and forth between the block diagram and biomolecular parameters are reported in Section 6 as a special case of (30) and (31).

## B Root Locus Analysis

To carry out a standard root locus analysis, the closed-loop transfer function should be rewritten in the following form

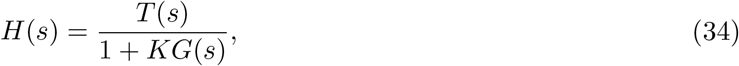

where *K* is the constant gain of interest (e.g. *K*_*I*_ or *K*_*P*_), such that 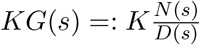 represents the loop gain, and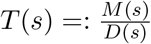 is a rational function of *s* which does not play a role in the closed-loop root locus. For the rAIF topology in Fig. 2(c) which realizes a filtered (I + FF) controller, (18) can be rewritten in the form of (34) as

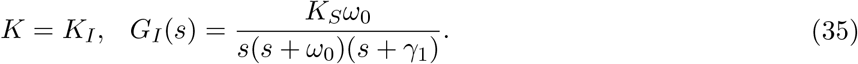

The root locus starts (at *K*_*I*_ = 0) from the poles (0, −*ω*_0_, −*γ*_1_) of *G*(*s*) and ends (at *K*_*I*_ → ∞) at its zeros (*s* → ∞ because *N* (*s*) = *K*_*S*_*ω*_0_ is a constant). As *K*_*I*_ is increased from zero, the first root-locus branch starting from the most negative open-loop pole, − max(*γ*_1_, *ω*_0_), moves on the real axis toward −∞. The other two branches move toward each other and break away from the real axis and approach two asymptotes intersecting with the real axis at −(*γ*_1_ + *ω*_0_)*/*3 with angles *π/*3 and −*π/*3. The break-away point of the root-locus branch starting from *s* = − min(*γ*_1_, *ω*_0_) and *s* = 0 is at

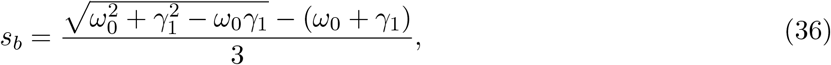

and so it is easy to show that 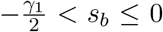. In fact, the fastest response which can be achieved by an infinite cutoff frequency *ω*_0_ is limited by a threshold dictated by 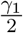.

## C Pole Placement

In this section, we derive the bounds on the achievable poles for the two negative actuation scenarios of sAIF: repression and degradation. Placing the three poles at a single location *s* = −*a*, allows us to express (*K*_*P*_, *K*_*I*_, *ω*_0_) in terms of the birth-death parameter *γ*_1_, the sensing gain *K*_*S*_, and the placed pole −*a* as shown in (20). Note that the supporting input *ū* is calculated using the equation *ū* − *γ*_1_*r* = 0, where *r* := *μ/θ* represents the set-point.

### C.1 Repression

Plugging (*K*_*P*_, *K*_*I*_, *ω*_0_) in the coverage condition in (13) yields

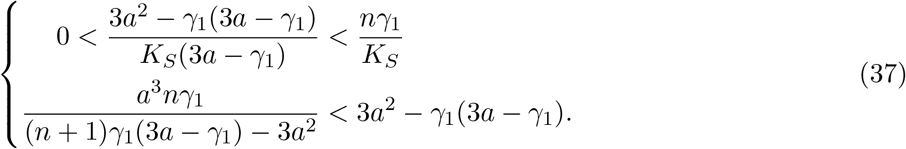

From the first inequality, we get

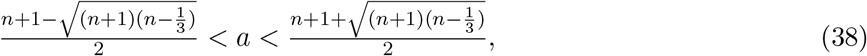

and from the second, we get *ζ*_1_ *< a < ζ*_2_ where *ζ*_1_, *ζ*_2_ are the two positive roots of the following fourth-order polynomial equation given by

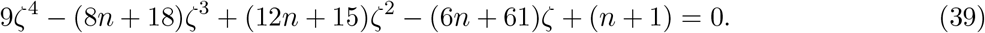

Calculating the intersection of the two inequalities yields the bounds for the achievable poles *s*_*l*_(*n*)*γ*_1_ *< a < s*_*u*_(*n*)*γ*_1_ where

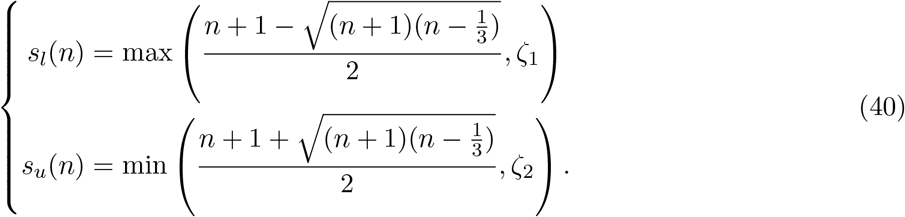

In the case of repression without cooperativity, it is not possible to place the three poles at a single location. However, we can still study the dynamics by placing the poles at two different locations instead of one. To this end, assume two poles are placed at *s* = −*a*_1_ and one pole at *s* = −*a*_2_. Equating the closed-loop characteristic polynomial in this case (*s* + *a*_1_)^2^(*s* + *a*_2_) to the denominator of *H*_s_(*s*) gives the expression of the PI gains (*K*_*P*_, *K*_*I*_) and the cutoff frequency *ω*_0_ in terms of the birth-death parameter *γ*_1_, the sensing gain *K*_*S*_ and the placed pole locations −*a*_1_, −*a*_2_ as

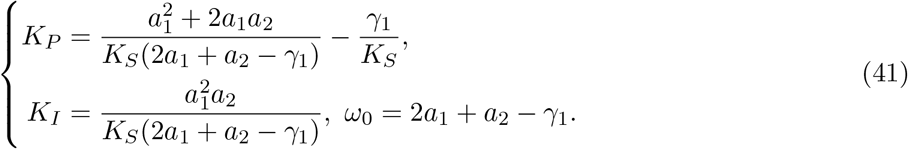

Plugging (*K*_*P*_, *K*_*I*_, *ω*_0_) in the coverage condition in (13) yields

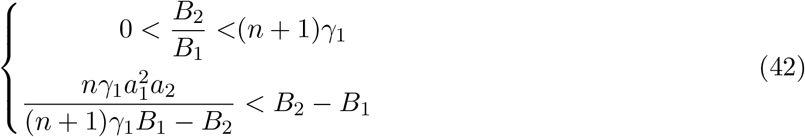

where *B*_1_ = 2*a*_1_ + *a*_2_ − *γ*_1_, 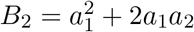. Rewriting *a*_1_ = *b*_1_*γ*_1_ and *a*_2_ = *b*_2_*γ*_2_, the inequalities in (42) simplify to

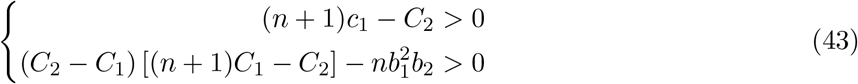

where *C*_1_ = 2*b*_1_ + *b*_2_ − 1, 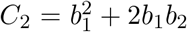. One can rely on graphical tools to calculate the intersection of the two inequalities as demonstrated in Fig. 4(d).

### C.2 Degradation

Plugging (*K*_*P*_, *K*_*I*_, *ω*_0_) in the coverage condition in (17) yields

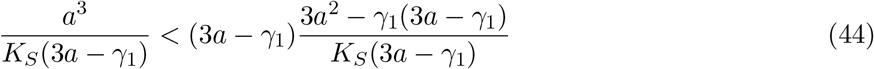

which simplifies to 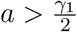.

